# Development and assessment of a new multichannel electrocutaneous device for non-invasive somatosensory stimulation for magnetic resonance applications

**DOI:** 10.1101/2024.05.27.595320

**Authors:** Carolina Travassos, Alexandre Sayal, Paulo Fonte, Nuno Carolino, Bruno Direito, Luis Lopes, Sonia Afonso, Tania Lopes, Teresa Sousa, Miguel Castelo-Branco

## Abstract

Electrocutaneous stimulation (ES) relies on the application of an electrical current flowing through the surface of the skin, eliciting a tactile percept. It can be applied for somatosensory mapping approaches at functional magnetic resonance imaging (fMRI) to obtain somatotopic maps illustrating the spatial patterns reflecting the functional organization of the primary somatosensory cortex (S1). However, its accessibility remains constrained, particularly in applications requiring multiple stimulation channels. Furthermore, the magnetic resonance (MR) environment poses several limitations in this regard. This study presents a prototype of a multichannel electrocutaneous stimulation device designed for somatosensory stimulation of the upper limbs of human participants in an MR environment in an inexpensive, safe, customizable, controlled, reproducible, and automated way. Our current-controlled, voltage-limited, stimulation device comprises 20 stimulation channels that can be individually configured to deliver various non-simultaneous combinations of personalized electrical pulses, depending on the subject, stimulation site, and stimulation paradigm. It can deliver a predefined electrical stimulus during fMRI acquisition, synchronized with the stimulation task design and triggered upon initiation of the acquisition sequence. Regarding device assessment, we conducted tests using an electrical circuit equivalent to the impedance of the human body and the electrode-skin interface to validate its feasibility. Then, we evaluated user acceptability by testing the device in human participants. Considering the stringent conditions of the MR environment, we performed a comprehensive set of safety and compatibility evaluations using a phantom. Lastly, we acquired structural and functional MR data from a participant during a somatosensory stimulation experiment to validate brain activity elicited by electric stimulation with our device. These assessments confirmed the device’s safety in fMRI studies and its ability to elicit brain activity in the expected brain areas. The scope of application of our device includes fMRI studies focused on somatosensory mapping and brain-computer interfaces related to somatosensory feedback.

## 1 INTRODUCTION

Neuroimaging research on somatosensory processing in the human brain is valuable for understanding its underlying neurobiology, mechanisms of disease and neuro-plasticity, and recovery after injuries to sensorimotor cortical areas (*e.g.*, due to stroke) or bodily injury (e.g., limb amputation) (1, 2). Non-invasive somatosensory stimulation benefits from mechatronic devices since they can provide controllable and reproducible stimuli. These properties are crucial for a more precise interpretation of brain dynamics, particularly within the somatosensory system, given the finely tuned functional properties of human skin tactile receptors (mechanoreceptors) (3, 4). In contrast, manual stimulation lacks consistent control over force, frequency, velocity, and coverage (5). The variability inherent to manual stimulation represents a significant limitation and hinders the generalization of findings (6, 7). Several actuation principles, including piezoelectric, pneumatic, electromagnetic, and electric, have been applied, each with relative advantages and disadvantages. The choice of an actuation principle depends on several factors, namely, available resources (materials, funding, and expertise) and the intended application(s) (9, 11).

The magnetic resonance (MR) environment imposes restrictions on the applicability of mechatronic stimulation devices due to the presence of a high magnetic field (static field), switching magnetic gradients, and radiofrequency (RF) pulses (8–11). Therefore, such devices must undergo rigorous safety and compatibility assessments before their use in the scanner room (9, 11).

To map the cortical representation of the upper limb in the human primary somatosensory cortex (S1) using functional magnetic resonance imaging (fMRI), we developed an electrical somatosensory stimulation device. Electrocutaneous stimulation (ES) involves the application of electric pulses to the skin using electrodes placed on the skin’s surface. This process activates skin receptors that process somatosensory information and elicit tactile-like sensations (12, 13). Previous studies have mapped somatosensory brain areas, investigating the impact of stimulation parameters on blood oxygen level-dependent (BOLD) fMRI response and somatosensory evoked potential studies (12, 14, 15). However, most ES devices are constrained by a limited number of stimulation channels, typically composed of two electrodes, thus limiting their range of applications (e.g., hand or median nerve stimulation).

The solution presented here is highly customizable, as it features twenty stimulation channels, portability, cost-effectiveness, and ease of development. Therefore, it can be employed in a broad range of MR/fMRI experiments. The present study describes the design, development, and assessment of a novel electrical somatosensory stimulation device. The evaluation process consisted of several steps. First, we tested the device’s ability to generate precise ES pulses using a circuit equivalent to the electrode/skin interface and internal body impedance. Then, we evaluated its ability to produce controllable and perceptible tactile sensations in a group of healthy participants. Following that, we conducted safety and compatibility assessments of the device in the MR environment using a phantom. Finally, we evaluated the device’s capability to generate stimuli for mapping the critical representation of the right upper limb in the S1 of a healthy human volunteer during MR imaging.

## 2. MATERIALS AND METHODS

An electrical stimulator intended for use in the MR environment must adhere to rigorous safety standards, encompassing both electrical safety and compatibility with the MR setting. In the following sections, we detail the design and functioning of our stimulation device, as well as how it aligns with the main standards established by international committees responsible for addressing these specific issues, following the recommendations of a recent systematic review on the field (11).

### 2.1. Overview of the somatosensory stimulation device

The device presented in this work was designed to stimulate the dominant upper limb of human participants during fMRI acquisitions for mapping its cortical representation in S1. ES is delivered through 20 independent stimulation channels under computer control, allowing for the predefinition of stimulation parameters, such as current, maximum voltage, and timing (pulse width and frequency and stimulation duration), according to the participant’s tactile perception and the experiment’s goals. Additionally, it enables the prior definition of the stimulation paradigm, including stimulation order, number of repetitions, and other customizable aspects.

In summary, the main stimulation device, housing the components responsible for generating the electrical stimulus, is controlled by both the MRI console computer, which receives the MR trigger and timely initiates the stimulation sequence using MATLAB R2019b, and by the Control interface computer that runs the Arduino C++ code (in Arduino IDE) responsible for managing the stimulation properties and experimental protocol. Considering the limitations of using ferromagnetic materials inside the scanner room, the main stimulation device was kept in the control room and connected to the penetration panel, employing RF filters. Within the scanner room, MR conditional^1^ cables and electrodes are responsible for delivering the stimulation signals to the participant. The switchboard controls the active and ground electrodes at each time according to the predefined stimulation sequence. The electric stimulus is monitored in real-time using a digital oscilloscope (Picoscope) and the control interface computer. Devices requiring power are connected to power sources, including an uninterrupted power supply (UPS). Figure 1 provides a schematic representation of the stimulation setup implemented in the MR environment showcasing the aforementioned components.

**Figure 1.**
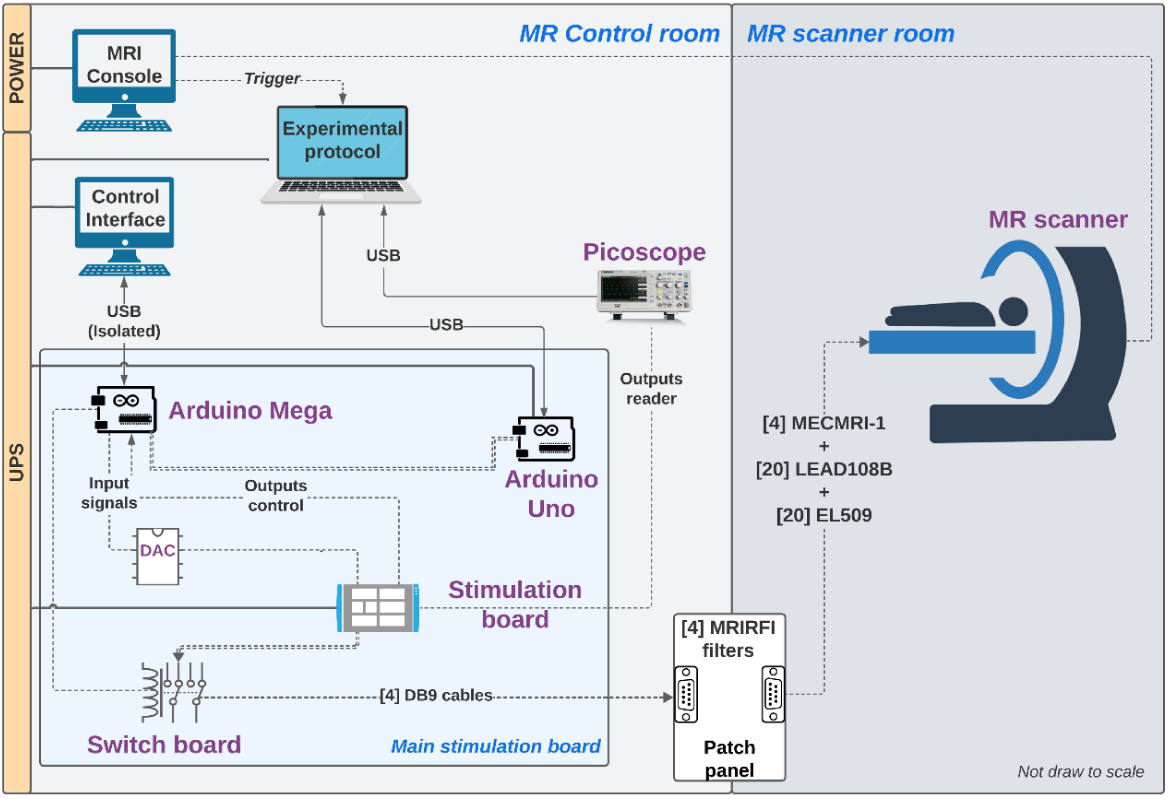
Stimulation setup with a participant in the magnetic resonance (MR) environment: schematic representation of the setup, including the main stimulation device (housing the components responsible for stimuli generation), the digital oscilloscope (Picoscope) and control interface computer (for quality assurance), the experimental protocol computer (to execute the experimental protocol), and the MR console computer (to receive the MR trigger at the start of the acquisition and enable synchronization with the stimulation protocol). The communication between the scanner and the control rooms is made through the path panel recurring to radiofrequency filters (MRIRFI). In the scanner room, are represented the scanner, participant, electrodes (EL509), and the respective cables (MECMRI and LEAD 108B). It also represented the power sources (power and uninterrupted power supply - UPS) and the remaining cabling.

#### 2.1.2. Main stimulation device specifications

The main stimulation device consists of a pulse generator board, a switchboard, a digital-analog converter (DAC), two Arduinos, and an auxiliary board (Figure 2). The pulse generator board produces a current-controlled, voltage-limited, rectangular stimulation signal, controlled by analog continuous input voltages and by one digital pulse that defines the pulse timing. The outputs from the pulse generator are current (Imon) and voltage (Vmon). The full electrical diagram and description can be found in supplementary material, section 1.

**Figure 2.**
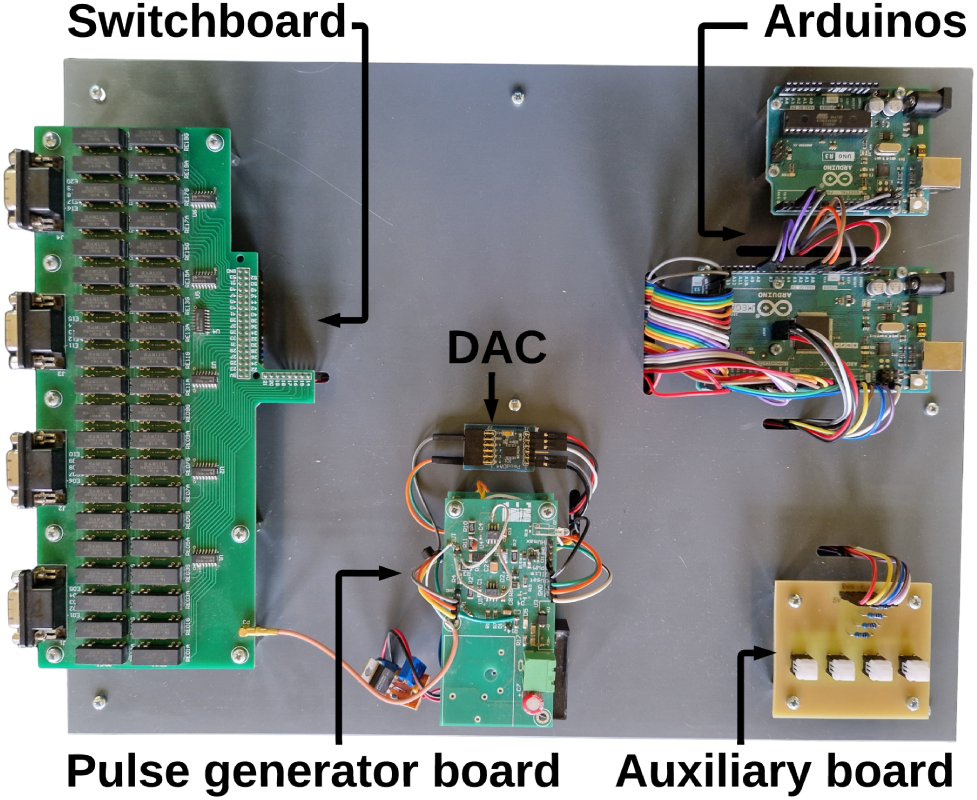
Photography of the main stimulation device, constituted by: Pulse generator board, Switchboard, Digital-Analog Converter (DAC), Arduinos Mega and Uno, and an Auxiliary board.

The pulse generator board is controlled by an Arduino Mega 2560 Rev (primary) and an Arduino Uno (secondary) (BCMI, New York, USA). The primary is responsible for defining the stimulation sequences and parameters (*i.e.*, order of stimulation) and is also used for quality control by displaying the outputs of the pulse generator board on the control interface computer. The secondary is responsible for receiving triggers from the MR scanner (*e.g.*, acquisition start) and sending this signal to the primary Arduino.

Communication between the primary Arduino and the pulse generator board is interfaced through a DAC (PmodDA4, Digilent Inc., Pullman, USA) using the serial peripheral interface (SPI) communication protocol. This DAC generates two control voltages (VSet and ILim) that define the output current and voltage of the stimulation.

The switchboard comprises 40 relays (HE721C0510, Littelfuse, Chicago, USA), two for each stimulation channel. It is responsible for connecting each electrode either to the pulse generator board or to the circuit ground, defining the active and passive electrodes in use at any given moment. Additionally, it incorporates four DB9 connectors to connect to the cable assemblies used in electrode-related cabling. The corresponding electrical diagram and description can be found in supplementary material, section S2.

Lastly, the system also includes an auxiliary board. It allows the manual interruption of the stimulation and controls the stimulation electrode for some applications *(e.g.*: perception thresholds update - please see the section 2.5.2 for details).

The stimulation board/signal generator was developed in general accordance with electric guidelines published at International Electrotechnical Commission (IEC) 60601-2-10 for basic safety and essential performance of nerve and muscle stimulators (16). The two main requirements are: 1) for stimulus pulse outputs, the maximum energy per pulse shall not exceed 300 mJ when applied to a load resistance of 500 ohms; 2) for stimulus pulse outputs, the maximum output voltage shall not exceed a peak value of 500 V when measured under open circuit conditions (11, 16). The voltage limitation feature of the pulse generator (settable pulse-by-pulse) prevents the application of unpleasant voltage levels if the load impedance suddenly increases because of electrode disconnections or defective electrodes/leads.

#### 2.1.3. Cables, electrodes, and RF filters

The interface between the main stimulation board and the participant comprises components that can safely enter the scanner room. These components are commercially available from BIOPAC (BIOPAC Systems Inc, USA) and were chosen for our setup due to their MR safety properties: stimulation electrodes (EL509) - MR conditional electrodes designed for electrodermal applications (17); electrode leads (LEAD108B) - MR Conditional short leads to connect electrodes to the main cabling (18); coaxial cables (MECMRI-1) - MR conditional shielded cables used inside the scanner room to connect electrode leads to the penetration panel (19); penetration panel filtering (MRIRFIF) - MR conditional RF filter installed at the penetration panel in the control room responsible for providing stable, low-impedance-earthed ground and prevent electromagnetic interference from external RF sources (19). We used a medical grade isolated USB cable (ISOUSB-Cable B, IFTOOLS GmbH, Germany) between the secondary Arduino and the MR trigger receiver computer.

To ensure optimal safety and quality of the setup, we prevent coupling the electric component of RF emissions with cables and eliminate the presence of conductive loops. Additionally, we employed twisted pair cables and positioned the electrodes in a straight line perpendicular to the axis of the magnetic field (9, 11, 28).

#### 2.1.4. Other components

All components requiring power to operate (computers, Arduinos, and pulse stimulus generator board) connect to an UPS (UPS 1000VA-4, Safire, Europe) that provides electrical surge protection.

We used a digital oscilloscope (Picoscope 5000 Series - Pico Technology, Eaton Socon, Cambridgeshire, England) to measure and record the outputs from the pulse generator board in real time for quality assurance.

### 2.2. Stimulation characteristics

The developed device is capable of generating positive rectangular stimulation signals, with maximum energy per pulse of 6 mJ, within a current range of 0 to 5 mA, a voltage up to 70 V, pulse widths between 0.2 and 5 ms, and frequencies up to 2.5 kHz for each of the 20 independent stimulation channels.

### 2.3. Software and code availability

The code we developed to control the stimulation device, according to our experimental protocol, is available at https://github.com/CIBIT-UC/electric-stim. It is organized following the steps for a complete stimulation session with an adult human volunteer. In each folder, we placed the Arduino code that controls and encodes the stimulation procedure, together with supporting code that generates the predefined sequences of electrodes, currents, and voltages. The Primary Arduino (stimulation control) is responsible for:

- definition of the stimulation sequence of electrodes, currents, and voltages;
- communication with the DAC to set the VSet and ILim input voltages;
- generation of pulse control via pulse width modulation (PWM);
- reading Vmon and Imon to check necessary voltage and provided current;
- control of the switchboard;
- waiting for a trigger from the secondary Arduino to start the stimulation protocol; Secondary Arduino (trigger) is responsible for:
- interfacing with MATLAB, receiving the trigger signal;
- sending a start signal to the primary Arduino;
- sending a stop signal in case of emergency.

### 2.4. Device assessment (Phantom)

We started by assessing the device’s ability to generate controllable electrical stimuli (as predicted in the experimental design) outside the MR environment. Then, we evaluated MR safety, including induced currents, and compatibility, including device performance and image quality.

#### 2.4.1. Feasibility to generate controllable electrical stimuli

The device’s capability to generate controllable electrical stimuli was tested according to the setup presented in Figure 3. An electrical model equivalent to the body (1 KΩ resistance) in series with the electrode/skin impedance (50 KΩ resistor in parallel with a 20 nF capacitor) was used (20). Electrical stimulation consisted of an intermittent paradigm involving 0.2 ms positive rectangular pulses at a frequency of 100 Hz for a duration of 4 s and a current of 2 mA.

**Figure 3.**
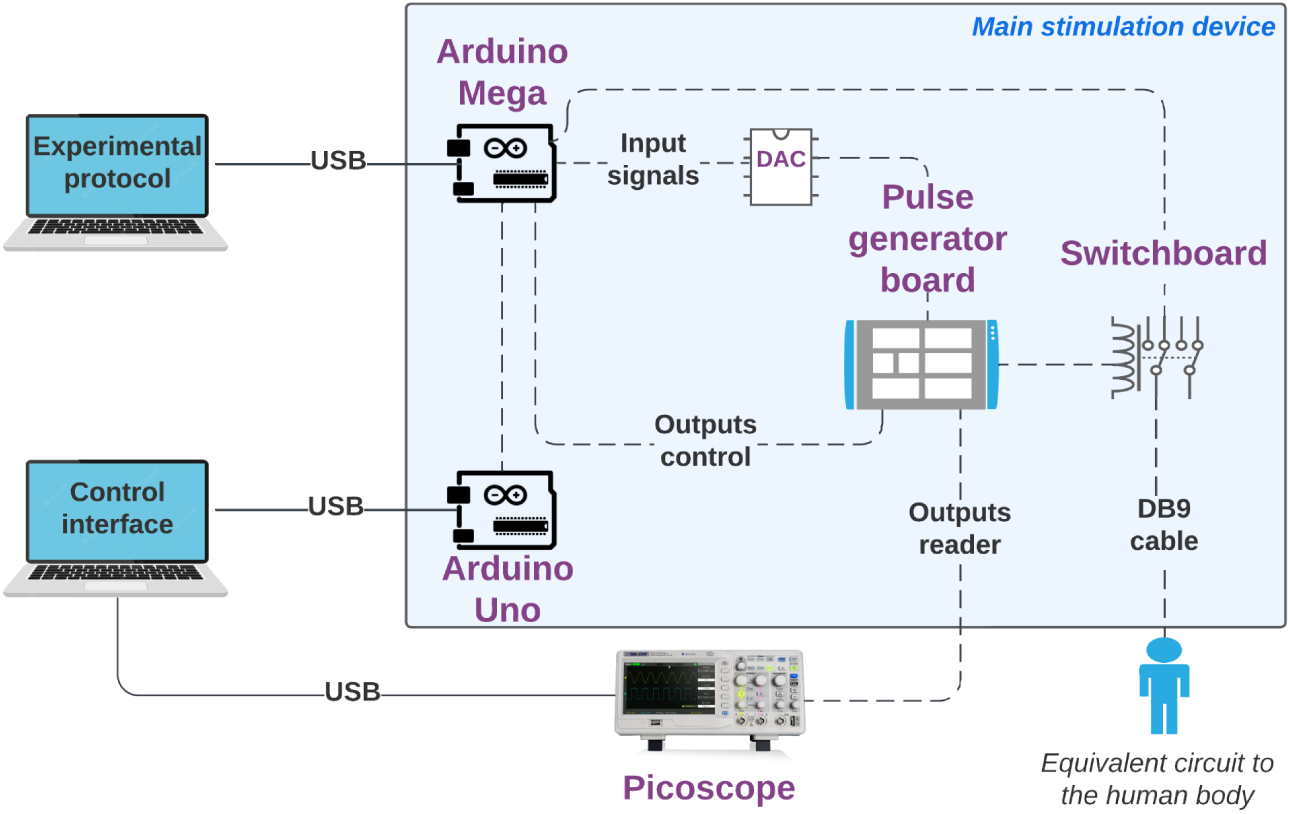
Schematic representation of the setup used to test the device’s feasibility in generating electrical stimuli. The setup includes a circuit simulating the impedance of the human body and the skin-electrode interface to evaluate the main stimulation device (represented by the blue box). A digital oscilloscope (Picoscope) and a control interface computer were employed for quality assurance, while another computer was used to execute the experimental protocol.

#### 2.4.2. Safety and compatibility in the MR environment

To verify that our device functions as expected in the MR environment and is not affected by RF pulse interference that could still exit the MR scanner room, we conducted safety and compatibility assessments that appeared most relevant in this context, following a recently proposed protocol (11).

MR assessments were performed on a 3 Tesla Magnetom Prisma fit scanner, equipped with a 64-channel head/neck coil, using a phantom (3.75 g NiSO4 x 6H2O, 5 g NaCl), all from Siemens Healthineers. Figure 4 illustrates the setup implemented (a), along with an image of the setup within the scanner room featuring the phantom (b).

**Figure 4.**
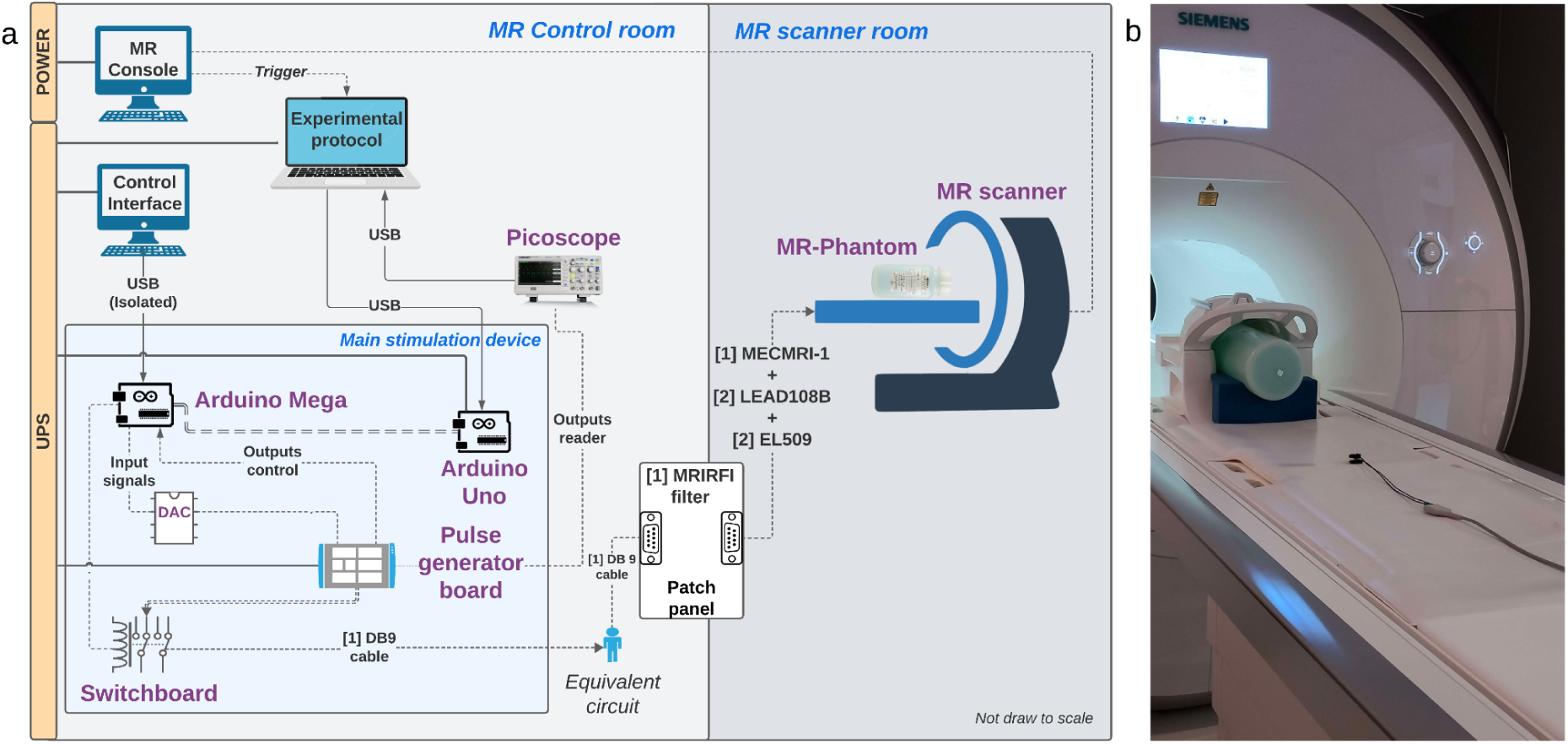
Stimulation setup with a phantom in the magnetic resonance (MR) environment. (a) Schematic representation of the setup, including the main stimulation device (blue box), the equivalent circuit (simulating the impedance of the human body and the electrode-skin interface), the digital oscilloscope and control interface computer (for quality assurance), the experimental protocol computer (to execute the experimental protocol), and the MR console computer (to receive the MR trigger at the start of the acquisition and enable synchronization with the stimulation protocol). (b) Picture of the phantom, electrodes, and cabling in the MR scanner room.

Functional images were acquired using a multi-band echo-planar imaging (EPI) sequence (27 contiguous slices, echo time: 37 ms, repetition time: 2000 ms, field of view: 186 mm, slice thickness: 1.6 mm, flip angle: 68°, GRAPPA = 2). Anatomical images were acquired using a high-resolution magnetization-prepared rapid acquisition gradient echo (MPRAGE) sequence (176 slices, echo time: 3.5 ms, repetition time: 2530 ms, field of view: 256 x 256 mm², flip angle: 7°, voxel size: 1 x 1 x 1 mm³).

Data acquisition consisted of four conditions: 1) no stimulation device in the setup (reference/baseline); 2) the device apparatus assembled but the device turned off (device off); 3) the device apparatus assembled and the device turned on (device on); and 4) the device operating (device operating). We tested both anatomic and functional acquisition sequences. Electrical stimulation consisted of an intermittent paradigm involving 0.2 ms pulses at a frequency of 100 Hz for a duration of 4 s. A current pulse of 1 mA was applied to one of the electrodes (electrodes’ leads/cabling were placed running through the MR table). Inter-stimulus intervals (ISI) of 6, 8, or 10 s were randomly applied between stimuli (mean = 6 s). The resistance between electrodes was approximately 1000 Ω, as measured with a digital multimeter.

##### 2.4.2.1. Safety

Constant monitoring of the stimulation outputs was performed in the control room using the digital oscilloscope and the outputs of the pulse generator board, displayed on the control interface computer. This setup allowed us to verify the stimulation outputs waveforms and assess the presence of any voltages induced by the electromagnetic environment of the scanner.

##### 2.4.2.2. Compatibility

Regarding compatibility assessment, we looked for qualitative and quantitative differences between the four image sets acquired, indicative of device compatibility with the MR scanner. Quantitative analysis was based on the following metrics: signal-to-noise ratio (SNR), temporal SNR (tSNR), and percentage image uniformity (PIU).

SNR compares the level of signal to the level of background noise. It was calculated based on the ratio between the signal correspondent to the average of a region-of-interest (ROI) encompassing approximately 75% of the phantom (ROI1) and the background noise corresponding to the standard deviation (SD) of four rectangular ROIs at the corners of the image (ROI2) (outside of the phantom and away from any artifacts) (Supplementary material, Figure S3.1); the SDs of the noise ROIs were averaged to obtain the mean noise and divided by the 0.66 Rician distribution correction factor to obtain total image noise (21). The SNR values were calculated for each slice of the anatomical scans and each slice and volume of the functional scans.

Additionally, for functional scans, we also calculated tSNR: a measure of image time course stability (22). tSNR was calculated by dividing the mean signal intensity of the voxel time series in ROI1 by the mean standard deviation of the voxel time course (mean of the SD values calculated for each volume separately).

PIU assesses the homogeneity of pixel values within the image; it was calculated based on thefollowing equation: 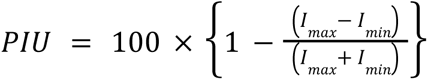, where *I_max_* and *I_min_* are the maximum and minimum intensity values extracted from ROI1, respectively (23). Then, the mean of the PIU values for each slice was calculated for anatomical scans, and the mean of the mean PIU values for each slice and volume was calculated for functional scans.

### 2.5. Generation of a perceptible tactile sensation, fMRI neural responses, and MR session workflow

We started by testing the device ability to generate perceptible tactile sensation in a human volunteer outside the MR environment. Then, we performed a pilot test in the MR environment following a mapping procedure to assess the device ability to elicit brain activity at the expected brain areas responsible for somatosensory processing. The study was approved by the local Ethics Committee and the participant signed the informed consent before taking part of it.

#### 2.5.1. Perceptible tactile sensation generation

The production of a perceptible tactile sensation was tested in a healthy participant (right-handed; female; 30 years) according to the setup presented in Figure 5. The test stimulus consisted of a pulse width of 0.2 ms, a frequency of 30 Hz, and lasted 4 s.

**Figure 5.**
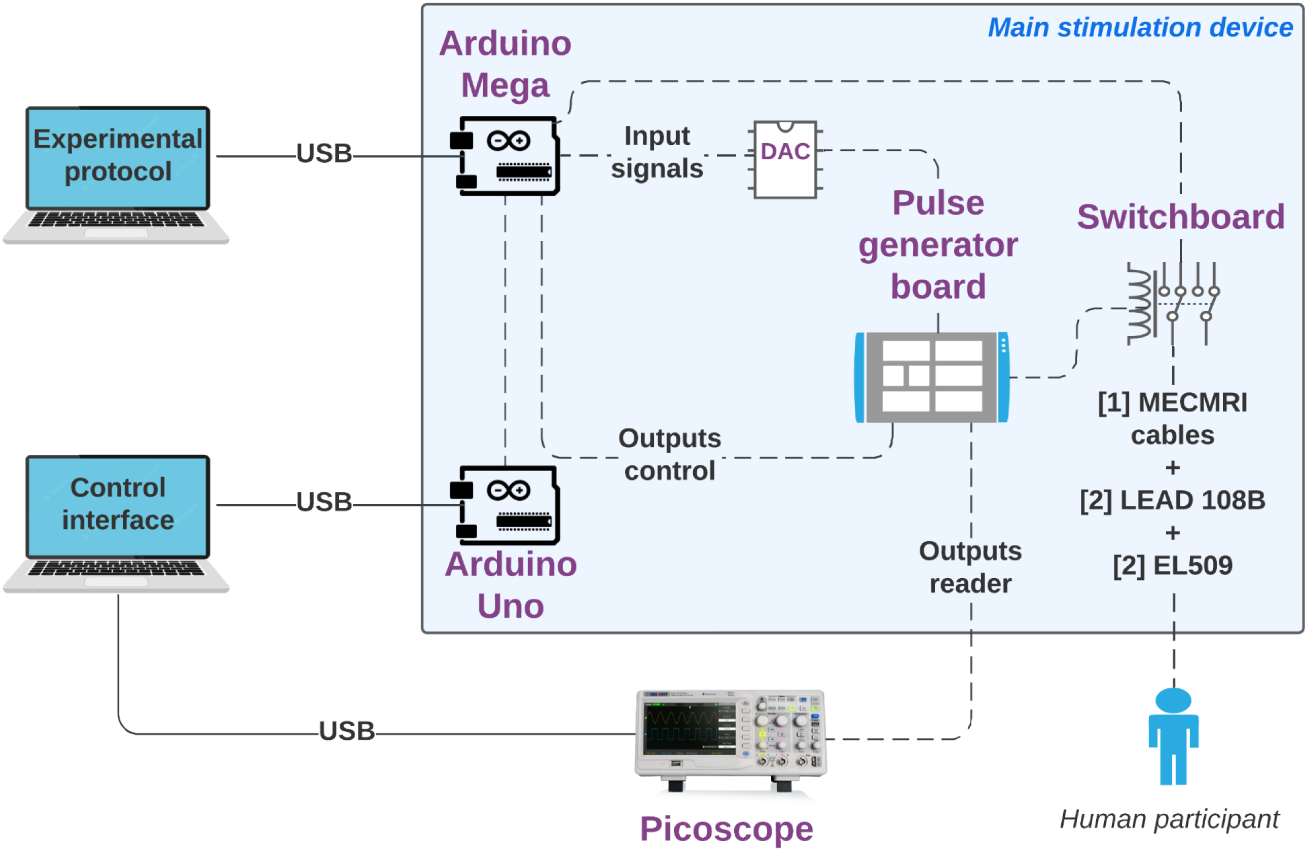
Schematic representation of the setup used to test the device’s feasibility in generating perceptible tactile sensation in a human participant. The setup includes the main stimulation device (represented by the blue box), a digital oscilloscope (Picoscope) and a control interface computer for quality assurance, and another computer to execute the experimental protocol.

#### 2.5.2. Pilot testing

To explore the capability of our device to elicit brain activity in the somatosensory cortex of a healthy participant (right-handed; female; 30 years) during an fMRI acquisition, we performed a pilot experiment. This experiment served three main purposes: 1) conducting a proof-of-concept imaging study to explore brain activation in regions linked to somatosensory function; 2) assessing the participant’s acceptance of the stimulation device in an MR environment; and 3) assessing the participant’s safe exit of the MR scanner in case of an emergency. Figure 1 illustrates the setup in the MR environment.

##### 2.5.2.1. Participant preparation

To prepare the skin for ES, we used abrasive pads (ELPAD) to remove non-conductive skin cells and ensure low contact impedance at the electrode attachment site. Additionally, we utilized electrode conductor gel (GEL101) (24, 25). Stimulation electrodes were placed on the dorsal side of the upper limb at designated stimulation sites. These sites are spaced proportionately based on the length of each upper limb segment; minimum spacing was determined based on literature about two-point discrimination tasks (26). Therefore, three electrodes were positioned on the middle finger, four on the hand, six on the forearm, and seven on the arm.

##### 2.5.2.2. Perception threshold determination

Perception threshold definition was performed based on a staircase procedure outside the MR environment. It involves the determination of the threshold current and voltage for each stimulation electrode according to a randomized order. The procedure consisted of applying 2 mA (duration: 4 s; frequency: 100 Hz; pulse width: 0.2 ms) and adjusting it in increments or decrements based on participant responses ("yes" or "no") to feeling the stimulation. The step sizes were 1 mA, 0.5 mA, 0.25 mA, and 0.1 mA through successive stimulations.

##### 2.5.2.3. MR session workflow

After positioning the participant at the MR scanner, the session started with the update of the current values of the perception thresholds defined outside the MR environment. Current is adjusted by 0.25 mA if the participant reports feeling the stimulation (to ensure consistent perception due to the variability inherent to perception thresholds) and by 0.5 mA if the participant does not feel the stimulation, until a positive report is obtained or the maximum current is reached. Then, we run through all electrodes again to update the voltage values and to guarantee that stimulation of each electrode is felt (a procedure we call “electrode check”). The MR session included acquiring a reference anatomical/structural image, followed by four functional scans. The entire session lasted approximately 1 hour and 30 minutes.

The stimulation protocol was similar to that previously applied in the phantom assessment, but the current was maintained at the updated threshold level. Each electrode was stimulated two times per run, in a total of four runs.

#### 2.5.4. MR data preprocessing and analysis

The acquired data was preprocessed using the fMRIPrep pipeline version 23.2.3 (described in supplementary material, section 4) (27). First-level analysis was performed using a General Linear Model (GLM) for each run. The design matrix included a predictor for each stimulation electrode and confound predictors for the six motion parameters. The contrast All Electrodes > Baseline (uncorrected threshold, *p* < 0.01) was used to identify the responding brain areas.

## 3 RESULTS

### 3.1. Feasibility and production of tactile sensation assessment

Figure 6 shows the stimulation outputs waveforms for current - Imon (purple), voltage - Vmon (blue), and the output stimulus - Out (yellow) obtained from the electrical simulation (SPICE) (a), the circuit equivalent to the human body (b), and the healthy participant (c). The examination of the stimulator’s performance in the two tested conditions indicated that the frequency, amplitude, and pulse width of the output stimulation signals broadly confirmed the simulated ones. The volunteer who tested the device with different stimulation parameters also reported its ability to produce tactile percept sensations.

**Figure 6.**
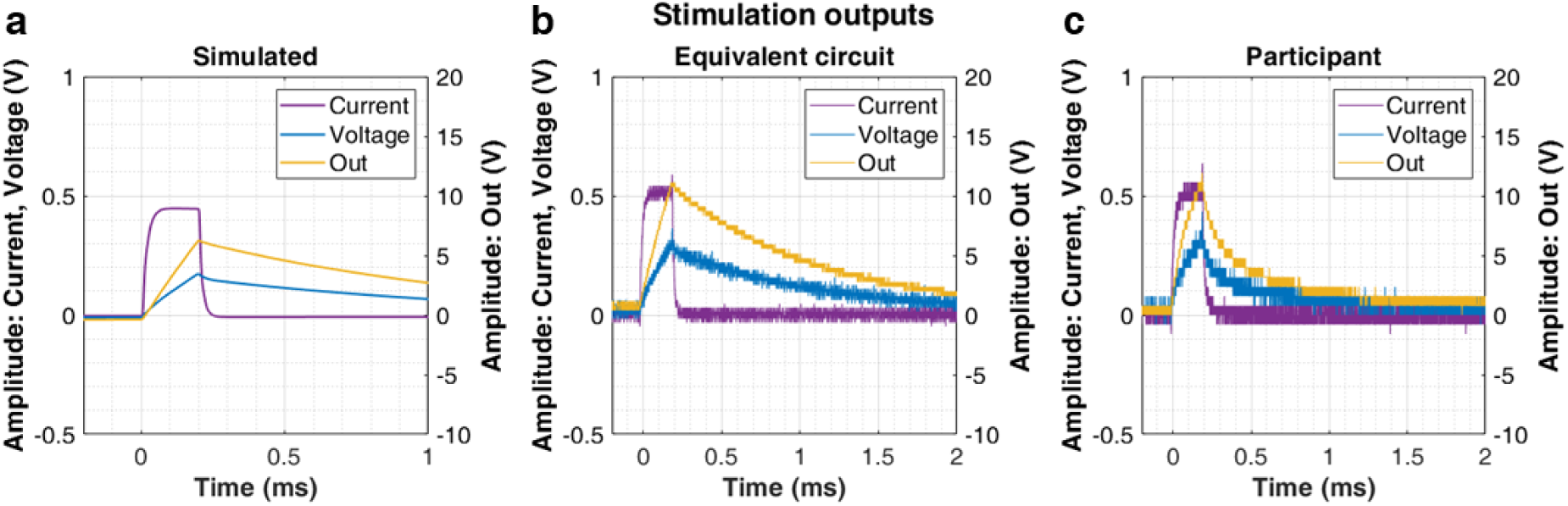
Stimulation outputs waveforms obtained outside the MR environment for a single stimulation pulse. Electrical simulation (SPICE) (a), the circuit equivalent to the human body (b), and the healthy participant (c). Stimulation performed with a frequency = 100 Hz, pulse length = 0.2 ms, and current = 1 mA (the actual voltage waveform depends on the impedance seen by the stimulator). Blue waveforms represent voltage (Vmon), purple represents current (Imon), and yellow represents the stimulus (Out).

### 3.2. Safety and compatibility in the MR environment with a phantom

Regarding safety assessment, we started by testing the existence of induced voltages in our setup for anatomical and functional acquisition sequences - Figure 7, a. Maximum noise amplitudes corresponding to the three main peaks that can be seen in the rightmost plot are on the order of 40 mV and 50 mV for anatomical and functional sequences, respectively, and therefore, negligible.

**Figure 7.**
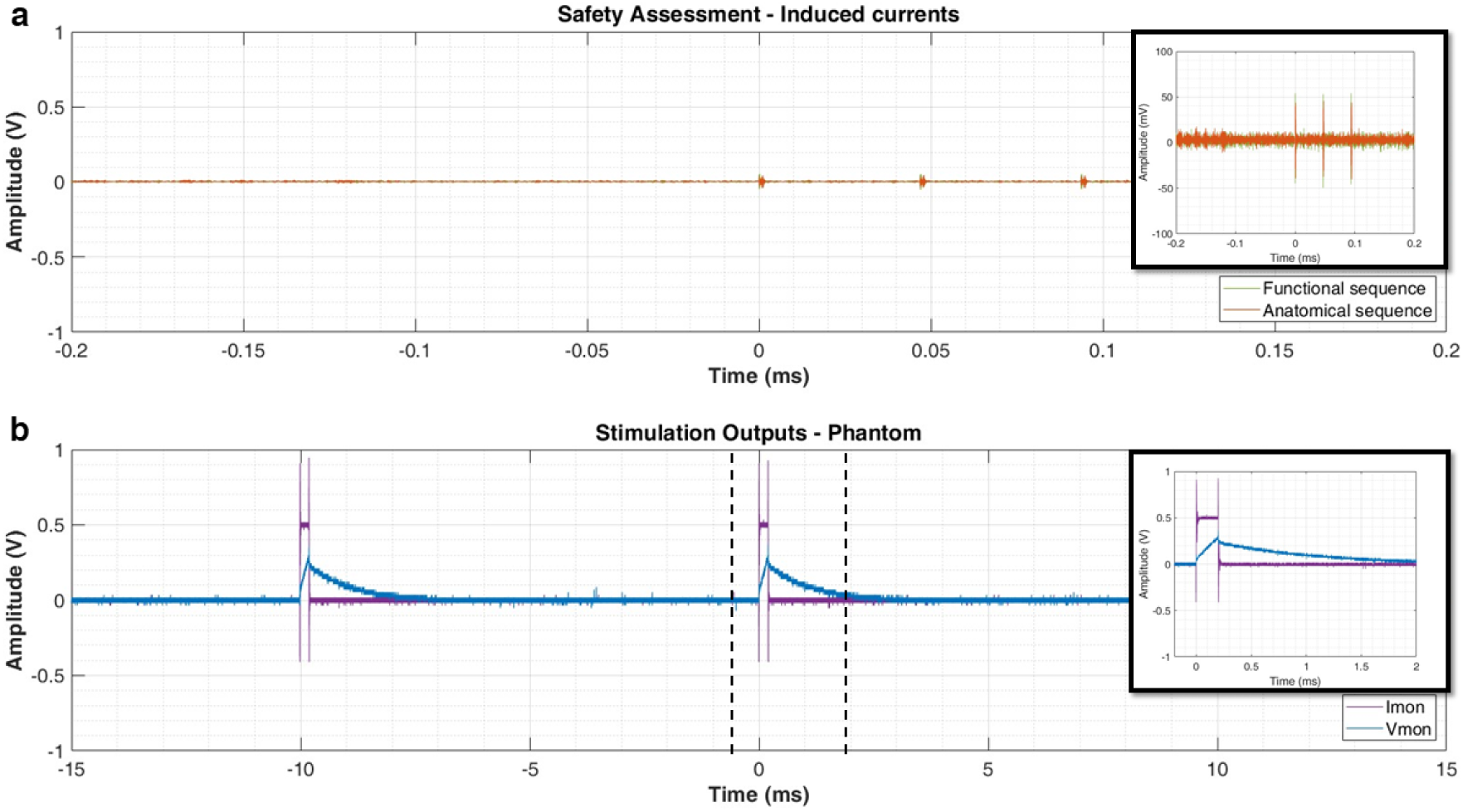
Stimulation output waveforms. (a) Induced voltage measurement during anatomical (orange) and functional (green) sequences; zooming of the amplitude axis for better visualization of noise in the mV scale. (b) Stimulation output waveforms of the functional acquisition with the phantom in the MR environment. In blue, we represent the Vmon voltage (1 V in this reading corresponds to 40 V in the output) and in purple, we represent ILim (0.5 V in this reading corresponds to 1 mA in the output). A single pulse waveform is highlighted on the right. Stimulation consists of positive rectangular pulses of 0.2 ms at a frequency of 100 Hz and a current of 1 mA.

We also obtained the stimulation output waveforms during an fMRI acquisition with the phantom (Figure 7, b). The imaging acquisition did not influence stimulation properties (frequency, amplitude, and pulse width), as we did not find differences when comparing with the stimulation output waveforms of the device outside the MR environment (stimulation outputs waveforms are present in Figure S5.1 in the supplementary material).

Concerning compatibility assessment, no degradation of the image quality was visually found for both anatomic and functional datasets (special attention was taken to the more common artifacts: spatial distortion, signal loss, or blurring of the images) (Figure S3.2 in the supplementary material presents a set of images corresponding to the four tested conditions for a single slice). Moreover, the quantitative analysis (results presented in Table 1) revealed that the variations between the four tested conditions were below 5% (deviations are presented in supplementary material, Table S3.1). Therefore, these metrics confirmed the insignificant impact of the device on image quality.

**Table 1.** Metrics used to evaluate image quality in anatomic and functional sequences for the four conditions tested: no stimulation device in the MR setup (Baseline); the device apparatus assembled but the device turned off (device off); the device apparatus assembled and the device turned on (device on); the device operating (device operating)^2^.

| Anatomic sequence |  |  |  |  |
| --- | --- | --- | --- | --- |
| (mean ± SD) | Baseline | Device OFF | Device ON | Device operating |
| SNR | 688.11 ± 38.98 | 679.41 ± 39.01 | 678.36 ± 35.88 | 673.95 ± 38.76 |
| Image Uniformity | 54.34 ± 2.24% | 54.34 ± 2.19% | 54.36 ± 2.23% | 54.44 ± 2.19% |
| Functional sequence |  |  |  |  |
| SNR | 22.47 ± 0.50 | 22.39 ± 0.49 | 22.39 ± 0.42 | 22.33 ± 0.45 |
| tSNR | 39.12 ± 0.12 | 38.74 ± 0.11 | 38.52 ± 0.12 | 38.50 ± 0.11 |
| Image Uniformity | 77.46 ± 1.83% | 74.09 ± 0.56% | 74.20 ± 0.50% | 74.08 ± 0.58% |

<sup>2</sup> SNR - signal-to-noise ratio; tSNR - temporal SNR; SD - standard deviation

### 3.3. Pilot testing in a human volunteer

Here, we present the first proof-of-concept neuroimaging data obtained with our electrical stimulation device in a healthy participant. Qualitative observations of fMRI data confirmed proper device functioning without causing spatial distortion, signal loss, or blurring of the images. Moreover, the stimulation output waveforms (Figure 8) remained as expected in terms of frequency, pulse width, and current (Figure S5.2 of supplementary material compares the plot of the stimulation outputs waveforms for this scenario with the ones obtained outside the MR environment). We verified the existence of a random noise pattern at the current waveform with no experimental meaning (to view the random noise pattern in the stimulation outputs waveforms please refer to Figure S5.3 in the supplementary material).

**Figure 8.**
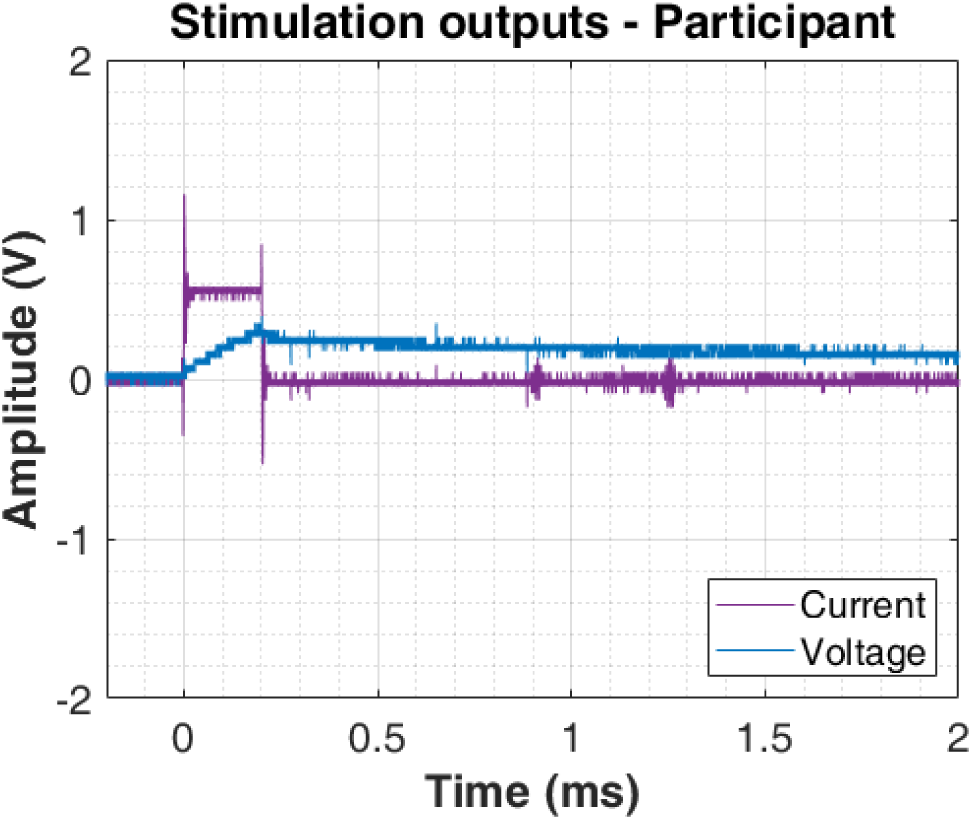
Stimulation output waveforms of a single stimulation pulse in the MR environment during a functional acquisition with a participant. Stimulation consists of positive rectangular current pulses of 0.2 ms at 100 Hz and a peak current of 1.10 mA. The purple waveform represents current (Imon) and the blue waveform represents voltage (Vmon).

The analysis of the fMRI data showed functional activation clusters in the somatosensory network, as shown in Figure 9. Brain areas pertaining to the sensorimotor network were identified (S1, Premotor cortex, Supplementary Motor Area, and others).

**Figure 9.**
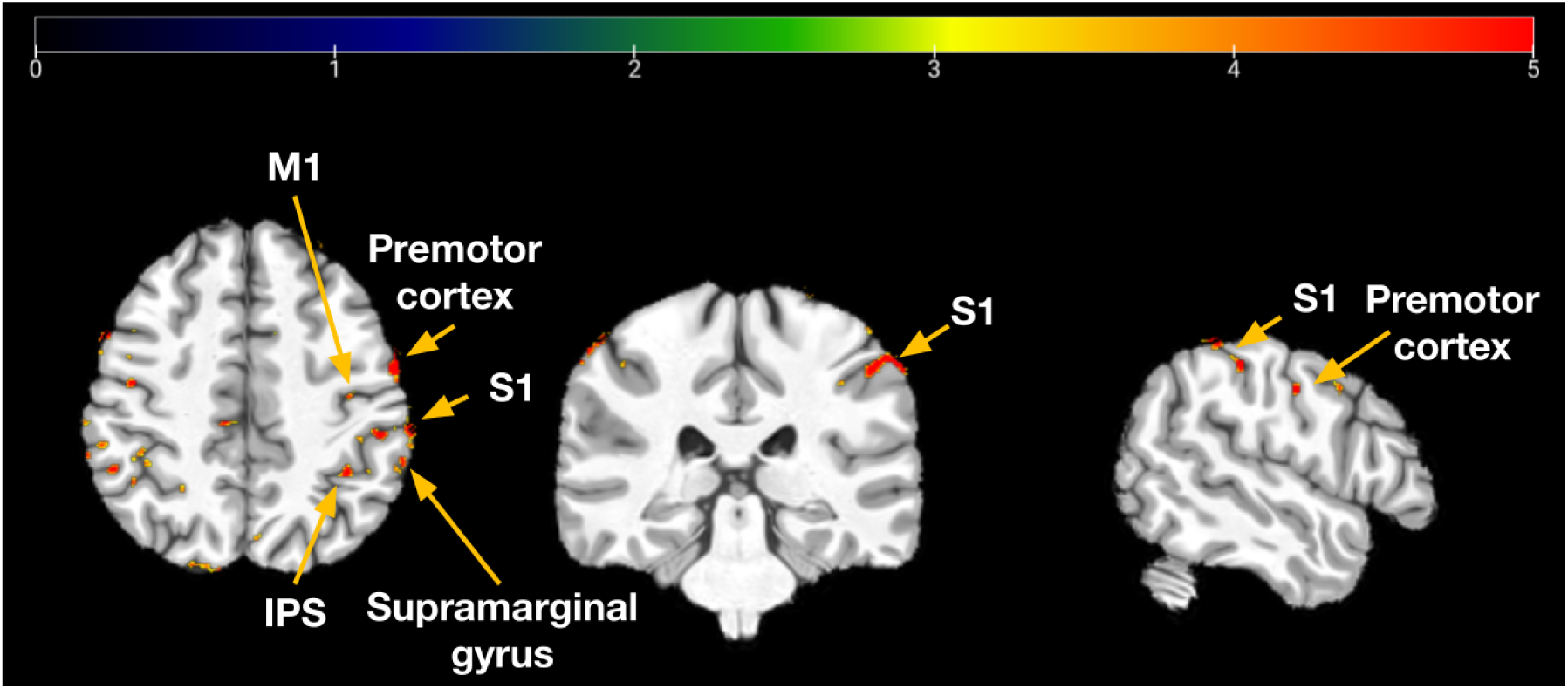
Statistical map resulting from the contrast “All Electrodes > Baseline” of all functional runs (uncorrected threshold, *p* < 0.01) from a healthy adult individual (MNI space). The color scale represents *t*-values. Brain areas identified: Primary Somatosensory Cortex (S1), Premotor Cortex, Primary Motor Cortex (M1), Supramarginal Gyrus, and Inferior Parietal Sulcus (IPS).

The participant did not report any adverse effect of stimulation nor additional discomfort caused by the presence of the stimulation setup. It was also verified that our setup does not hinder the safe exit of the participants from the scanner in case of an emergency since no component is fixed to the scanner bed.

## 4 DISCUSSION

The stimulation device presented here enables the generation of a predefined electrical signal to be delivered to a participant undergoing MRI/fMRI acquisition. It can also be used combined with other neuroimaging techniques. This device can produce a highly localized vibrotactile stimulus with a frequency range of 1–2.5 kHz and a maximum current and voltage of 5 mA and 70 V, respectively. Besides, it can synchronize the stimulation with a predefined task design and deliver stimulation at precise timing. These characteristics are of utmost importance for understanding the time course of brain activity related to somatosensory processing.

Our custom-made device was first assessed outside the MR environment, confirming its feasibility regarding stimulation outputs waveforms and producing perceptible tactile sensations. The implementation in the MR environment was designed to avoid potential sources of image degradation and interference with the device’s correct functioning. Negligible artifacts were characterized, confirming its safety and compatibility.

Several aspects contributed to the efficiency of the developed device. First, the coupling of the electric component of the RF emission with cables was avoided by ensuring they did not have resonant lengths (*i.e.*, lengths equal to half the wavelength of the RF field or its multiples). This issue, known as the antenna effect, occurs when certain components act like antennas, picking up electromagnetic interference or RF signals, causing image artifacts and heating (9, 11, 28). We also avoided conductive loops due to potential resonance heating, *i.e.*, increasing the ability of the loop to concentrate current, leading to excessive heating (9, 11, 28). To this end, we placed the cables straight in the *z*-direction of the main magnetic field of the scanner, in the center of the magnetic core and avoided direct contact between the cables and the participant. We also twisted the pair cables to reduce the conductive loop cross-sectional area and, this way, reduce the coupling between switching gradients (9, 11, 28). Last, RF filters at the penetration panel provided stable, low-impedance-earthed ground and prevented electromagnetic interference from external RF sources (9, 11, 28).

However, during fMRI acquisition with the participant, the stimulation current waveform (Imon) revealed slightly more induced noise than with the phantom. This could be due to the higher number of electrodes (and, therefore, leads and cables). The slightly increased spacing between them (in phantom assessments, electrodes were in contact with each other) likely increased the conductive loop cross-sectional area. Nevertheless, the superimposed induced current was negligible compared with the stimulation current.

The design of the MR implementation of the stimulation system presented here also took into account the optimal electrode position for electrical safety. As a main precaution, the stimulation electrodes were placed in a straight line perpendicular to the magnetic field axis (29, 30). Another important feature of our device is that it does not block participants from safely leaving the MR machine, in case of need, since it is not fixed to the scanner bed. This feature is of utmost importance due to the claustrophobic feeling often associated with MR scanners.

Lastly, the proof-of-concept imaging results demonstrate that our device effectively triggers brain activation in the expected brain regions, allowing for detailed somatotopic mapping (12). These findings demonstrate the effectiveness of our custom-made electrical stimulation device and experimental protocol in triggering activity in the expected brain areas, thereby validating its practical application.

### Limitations

According to IEC 60601-2, for electrical safety reasons, it is recommended to use batteries as the power source for this type of device (31). However, our device is connected to the main power supply via a UPS with surge protection, which is suboptimal. Moreover, our device can deliver a maximum current of 5 mA and a voltage of up to 70 V. Although this poses no limitations for the specific purpose outlined in this study, it could be a drawback for other applications, especially those involving pain assessment, which may need higher intensity levels. Nevertheless, this can be easily accommodated in an updated version of its design, as the main electronic limitation is the maximum voltage level, not the current.

## Conclusion

The device presented here enables safe, customizable, controlled, reproducible, and automated delivery of electric stimuli over 20 channels in an inexpensive manner. Moreover, it can deliver a predefined electrical stimulus during fMRI acquisition, synchronized with the stimulation task design and triggered upon initiation of the acquisition sequence. The modular assembly of our device enables its use in fMRI experiments without safety and compatibility issues. The system was designed for somatosensory mapping but can also be used for other applications. It can be in particular useful for human-machine interfaces including somatosensory feedback, such as prostheses control, sensory substitution, and sensory/sensorimotor restoration. Additionally, it could find use in pain neuroscience experiments and assessment of brain reorganization after lesions. Furthermore, this device offers an opportunity to enhance our understanding of electrotactile perception, a phenomenon that remains not fully understood.

## Footnotes

1 MR Conditional - MR safety label means that the item may safely enter the MR scanner room only under specific conditions provided in the labeling.

