## Supplementary Material for "Development and assessment of a new multichannel electrocutaneous device for non-invasive somatosensory stimulation for magnetic resonance applications"

### Section 1. Pulse generator board - circuit description

The schematic diagram of the pulse generator is shown in Figure S1. It generates an almost rectangular current-controlled positive pulse that is also limited to a maximum voltage. The circuit is based on the high voltage op amp LTC6090 and on the XPpower G01 DC/HVDC converter, with a maximum voltage of 100 V.

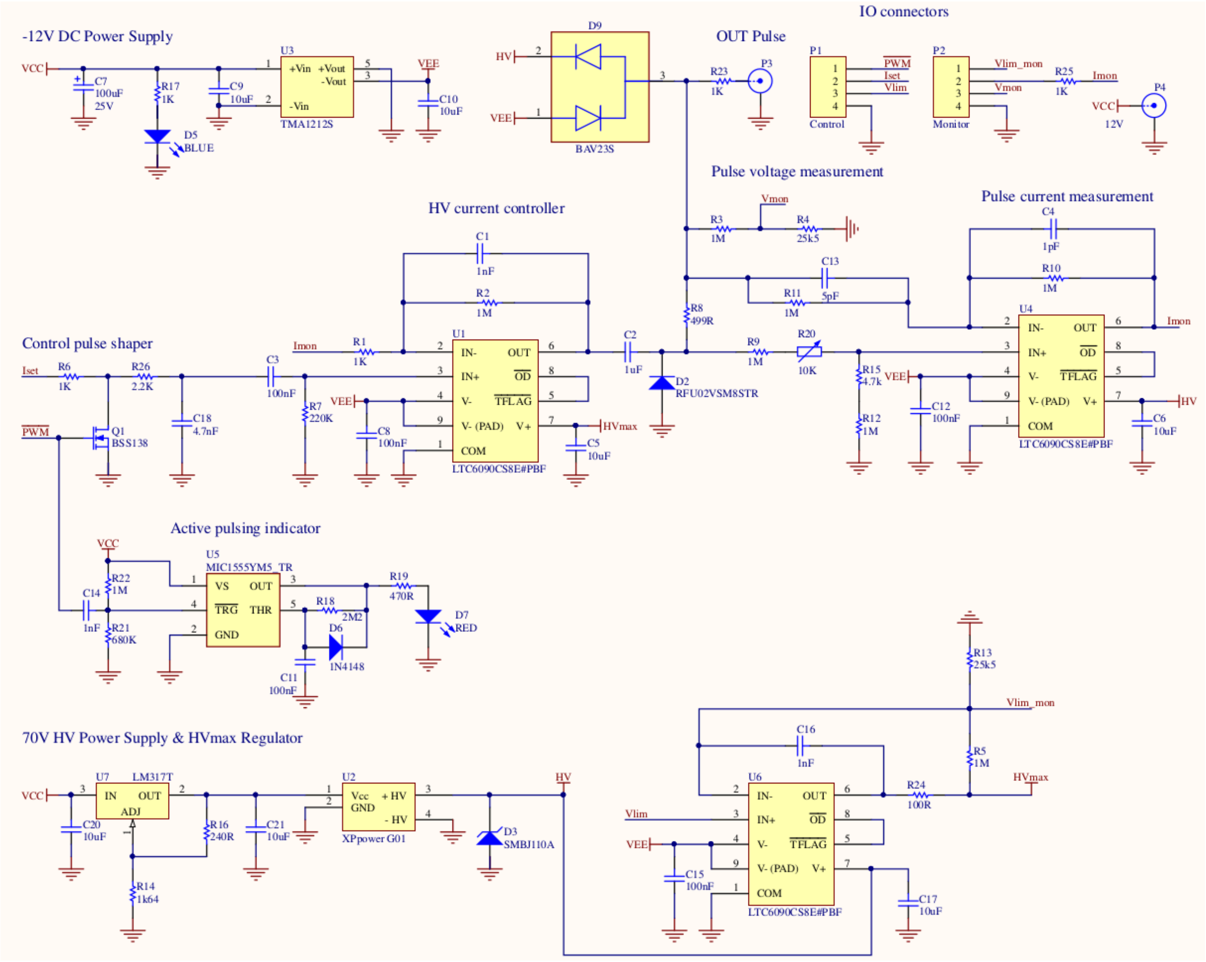


Figure S1.1 - Schematic diagram of the voltage-limited current-controlled pulse generator.

The circuit is analogically controlled by the input voltages entering the P1 connector: *Iset*, which defines the pulse height at a ratio of 2 mA/V; *Vlim*, which defines the maximum pulse voltage at a ratio of 40V/V; $\overline{PWM}$, which is a logical LVTTL-compatible active-low pulse that defines the pulse width and frequency.

Monitoring outputs of the pulse current and voltage waveforms, *Vmon* and *Imon*, with the same ratios as *Vlim* and *Iset*, are available for external measurement at the P2 connector. On the same connector the *Vlim_mon* output monitors the internal HV level (*HVmax*) set by *Vlim*, with the same voltage ratio.

The circuit is powered by a 12V medical grade AC/DC converter (GSM36E12-P1J, MW, Taiwan) at P4 (*VCC*) and it generates internally a -12 V power rail (*VEE*).

A 70 V fixed high voltage (*HV*) is generated by the proportional (100/12 V/V) G01 DC/HVDC converter that is fed at 8.4 V by a LM317 regulator. This defines the normal maximum possible pulse voltage and the TVS diode SMBJ110A hard-limits it to 110 V in the worst case. This voltage is then regulated by the U6 circuit to the value defined by *Vlim*.

The *control pulse shaper* circuit generates a positive rectangular voltage pulse with smooth edges that corresponds to the *Ilim* and $\overline{PWM}$ controls. For convenience and safety there is a visual indicator (LED) when in an active pulsing state.

This pulse is AC-coupled to the *HV current controller* circuit (U1) that is a proportional feedback controller with gain 1000 that compares the control pulse with the *Imon* output, forcing the pulse current to conform to the required shape and amplitude. This stage is powered by *HVmax*, limiting the pulse voltage to this value.

The HV pulse thus generated is coupled to the output stage via the 1 µF C2 capacitor. This is an important safety feature because in the worst case of total circuit failure it limits the energy of the pulse to $\frac{1}{2}1\mu F\,(110V)^2 = 6.0\,mJ$ , well within the safety regulations. The D2 fast diode provides the current return path for the pulse descending edge, rectifying the output signal.

The output stage comprises the current measuring resistor R8 = 499 Ω, which is sensed by a unity gain difference amplifier (U6) that generates *Imon*. The CMRR of the amplifier can be optimized by adjusting R20 with the output disconnected. There is also a voltage divider that generates *Vmon*. These circuits connect to the output only via 1 MΩ resistors so that they are unable to inject any significant current on the patient.

The BAV23s schottky diodes along with R23 protect the circuit from any spurious back-voltages that may appear at its output (P3).

The circuit can generate pulses from 0.2 ms to 5 ms at duty cycles up to 50%. Some example oscillograms taken from the Imon and Vmon outputs can be seen in Figure S1.2. In all cases the load was a model of the electrode-electrode impedance composed of 50 kΩ in parallel with 20 nF, both in series with 1 kΩ for the body internal resistance (20).

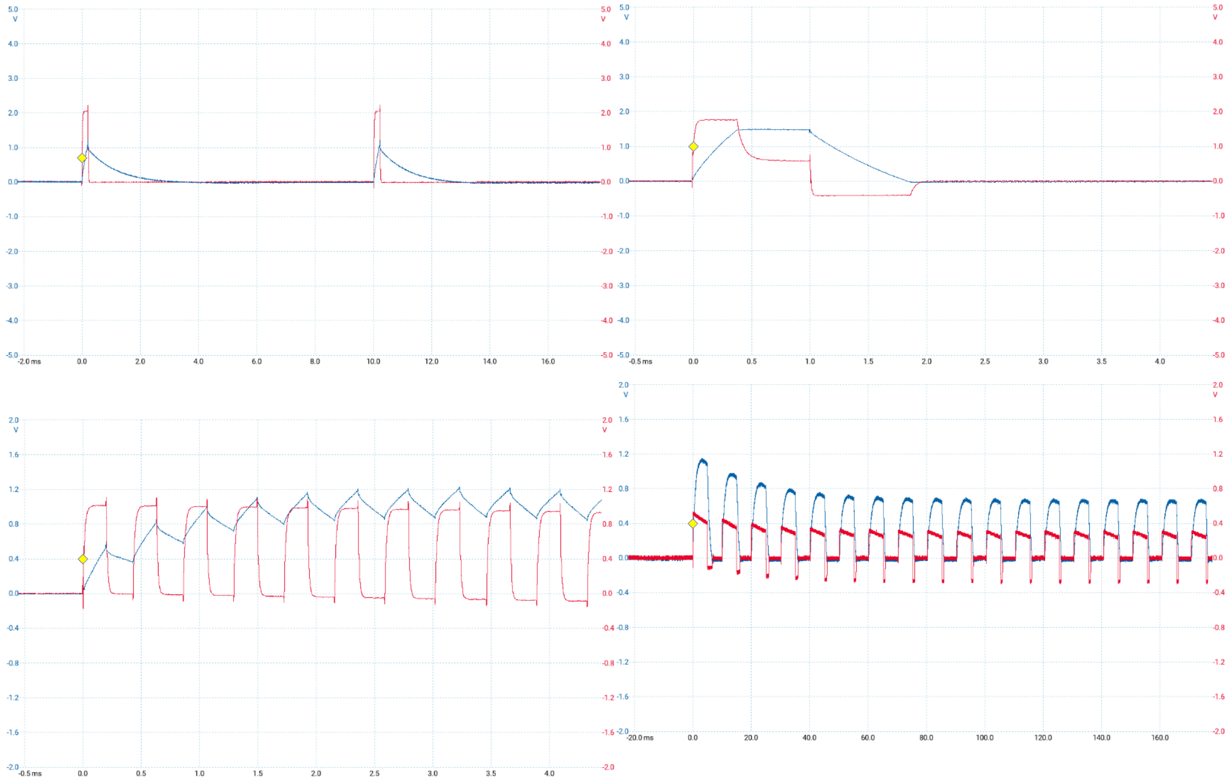

Figure S1.2 - Example oscillograms from the Vmon (blue) and Imon (red) outputs. Upper left: pulse width of 0.2 ms, at a frequency of 100 Hz, and current of 4 mA. Upper right: action of the voltage limitation feature set at 50 V. Lower right: pulse width of 0.2 ms, at a current of 2 mA, and 50% duty cycle. Lower left: pulse width of 5 ms, at a current of 1 mA, and 50% duty cycle; some pulse distortion on the order of 20% can be seen at this longer pulse width.

### Section 2. Switchboard - circuit description

The output of the pulse generator board is connected to the participant via a 20-channel multiplexer (“switchboard”) - Figure S2, top. The switchboard is constituted of 40 relays (two relays to control each electrode) that provides 3-state outputs: high-impedance (default), grounded or pulsed - Figure S2, bottom. This is based on Reed relays controlled one-to-one from the arduino I/O via ULN2003 buffers.

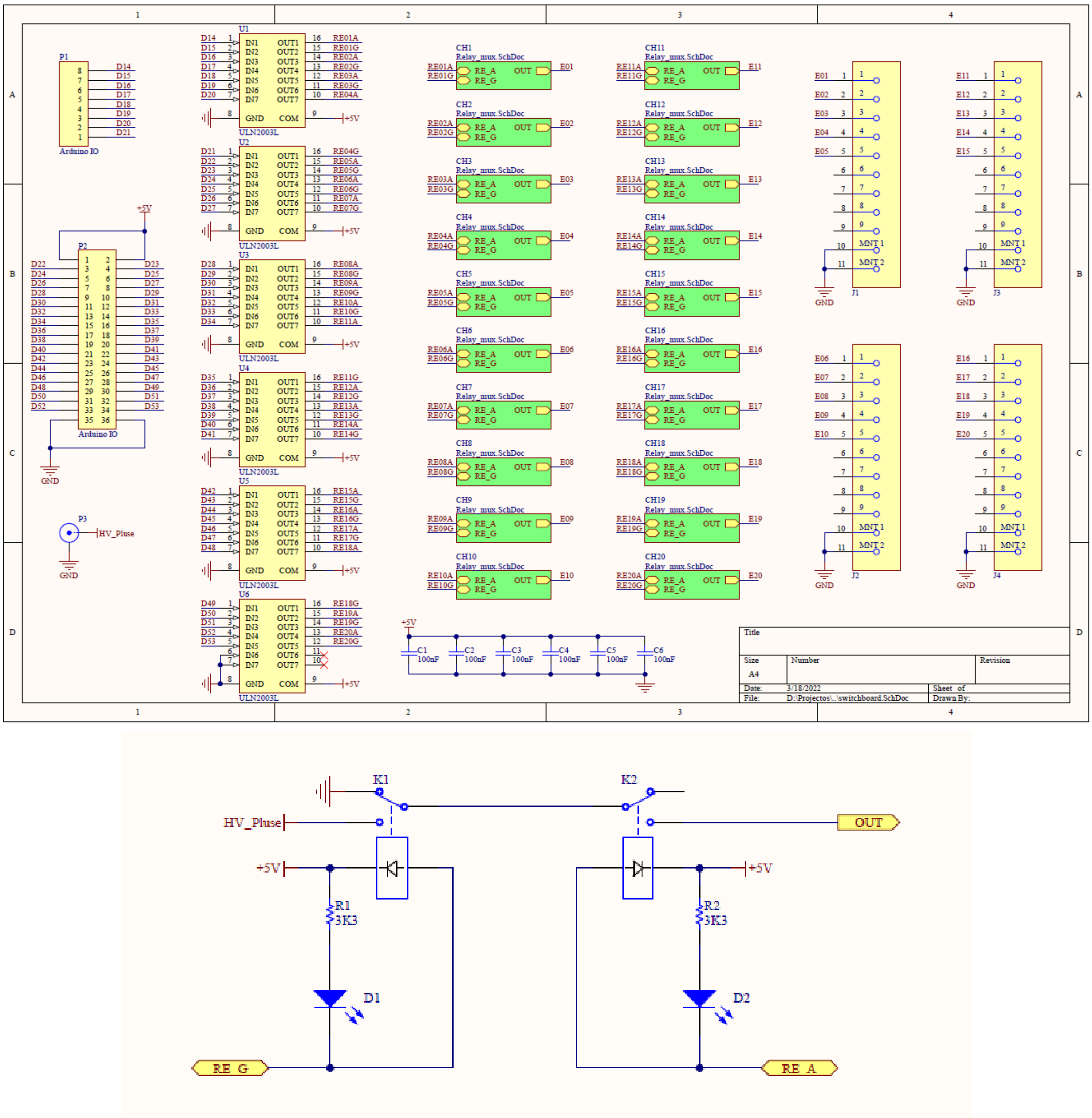


Figure S2 - Schematic diagram of the switchboard: Top) control circuit; Bottom) output stage.

### Section 3. Compatibility assessment

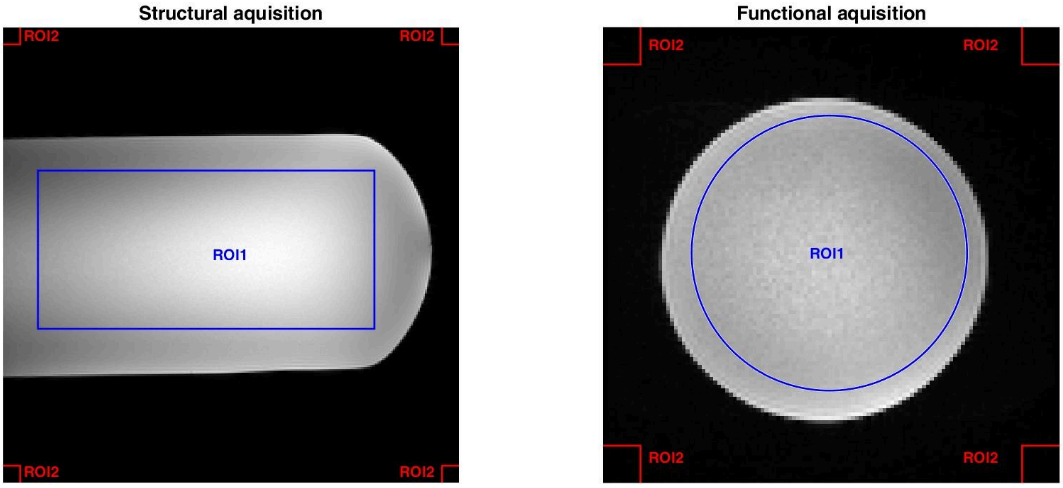


Figure S3.1. Schematic representation of the regions of interest (ROI) used to calculate signal-to-noise ratio at structural (left) and functional (right) data (one slice represented). ROI1 (blue) encompasses approximately 75% of the phantom (anatomical: 40 x 160 pixels; functional: 3853 pixels (radius = 35)) and was used to calculate the mean signal; ROI2 (red) corresponds to 10 x 10 pixels squares at the corners of the image and were used to calculate the mean standard deviation. Only slices including phantom were considered in the structural acquisition.

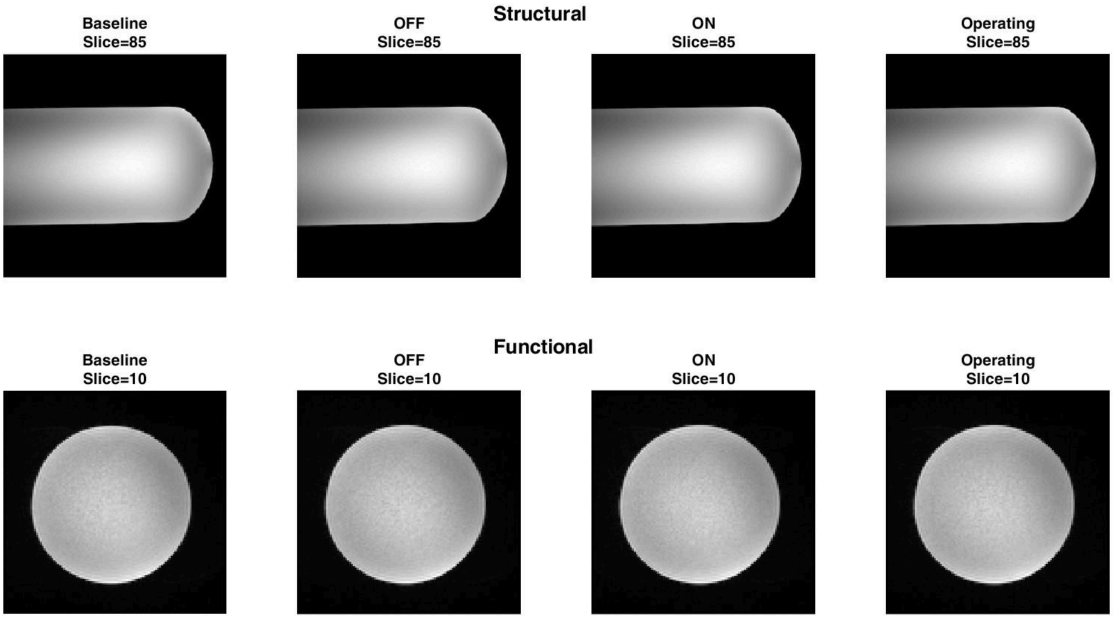


Figure S3.2. Comparison of the four conditions acquired in anatomic/structural (top) and functional (bottom) acquisitions with the phantom for qualitative image quality assessment.

Table S3.1. Deviations between the conditions tested (Device OFF, device ON, and device Operating) and the baseline condition based on the formula: $Deviation\ (\%) = \frac{metric_{condition} - metric_{baseline}}{metric_{baseline}} \times 100.$

| Anatomic sequence | | | |
|---|---|---|---|
| | Baseline vs Device OFF | Baseline vs Device ON | Baseline vs Device Operating |
| **SNR** | -1.26% | -1.42% | -2.06% |
| **Image Uniformity** | 0.00% | 0.05% | 0.19% |
| **Functional sequence** | | | |
| **SNR** | -0.37% | -0.33% | -0.64% |
| **tSNR** | -0.96% | -1.52% | -1.58% |
| **Image Uniformity** | -4.36% | -4.22% | -4.37% |

### Section 4. fMRIPrep pipeline

T1-weighted (T1w) image was corrected for intensity non-uniformity with "N4BiasFieldCorrection" (from ANTs) and used as T1w-reference throughout the workflow. The T1w-reference was then skull-stripped with a Nipype implementation of the "antsBrainExtraction.sh" workflow (from ANTs). Brain tissue segmentation of cerebrospinal fluid, white matter, and gray matter was performed on the brain-extracted T1w using "fast" (from FSL). Brain surfaces were reconstructed using "recon-all" (from FreeSurfer). Lastly, spatial normalization to standard space (MNI152NLin2009cAsym) was performed through nonlinear registration with "antsRegistration" (from ANTs).

Functional data preprocessing included head-motion correction (using "mcflirt", from FSL), slice-time correction (using "3dTshift" from AFNI), susceptibility distortion correction, co-registration with the T1w-reference (using "bbregister", from FreeSurfer, which implements boundary-based registration), and normalization into Montreal National Institute

(MNI) standard space (using "antsApplyTransforms", from ANTs). No smoothing was applied.

### Section 5. Comparison of the stimulation outputs outside vs inside the MR environment

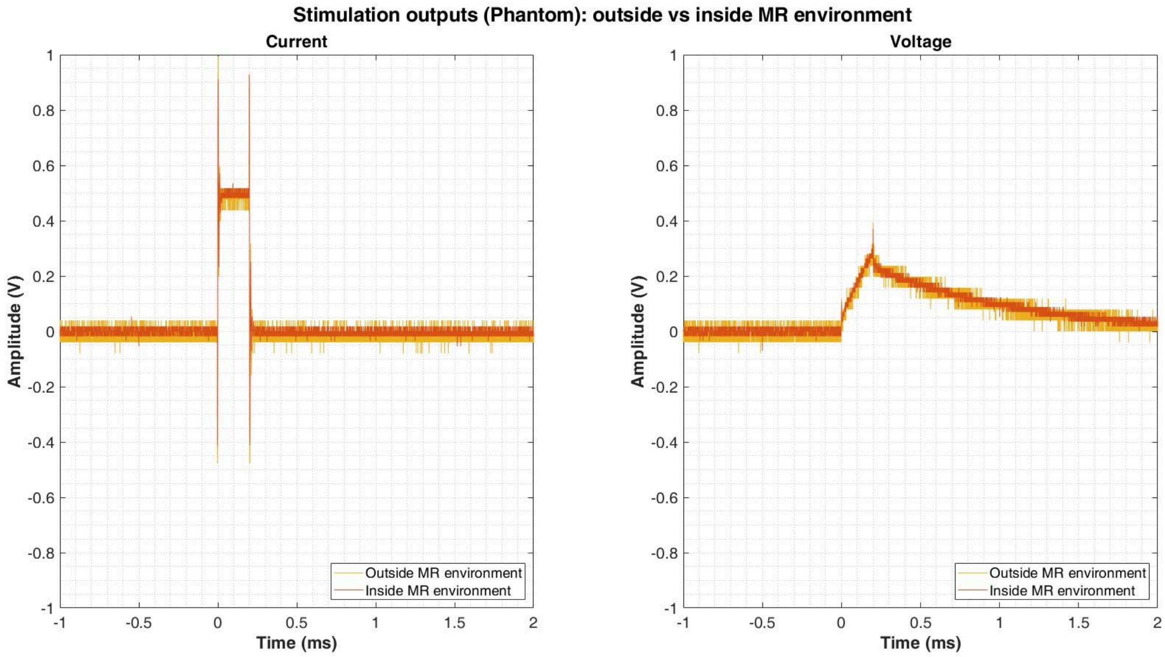


Figure S5.1 - Comparison of the current (left) and voltage (right) outputs measured inside (orange) and outside (yellow) the magnetic resonance (MR) environment for a single stimulus in the experiment with the circuit equivalent to the human body. The overlapping waveforms emphasize their similarity, indicating that the MR environment did not influence the performance of the stimulation device. Stimulation consists of positive rectangular pulses of 0.2 ms at a frequency of 100 Hz and a current amplitude of 1 mA.

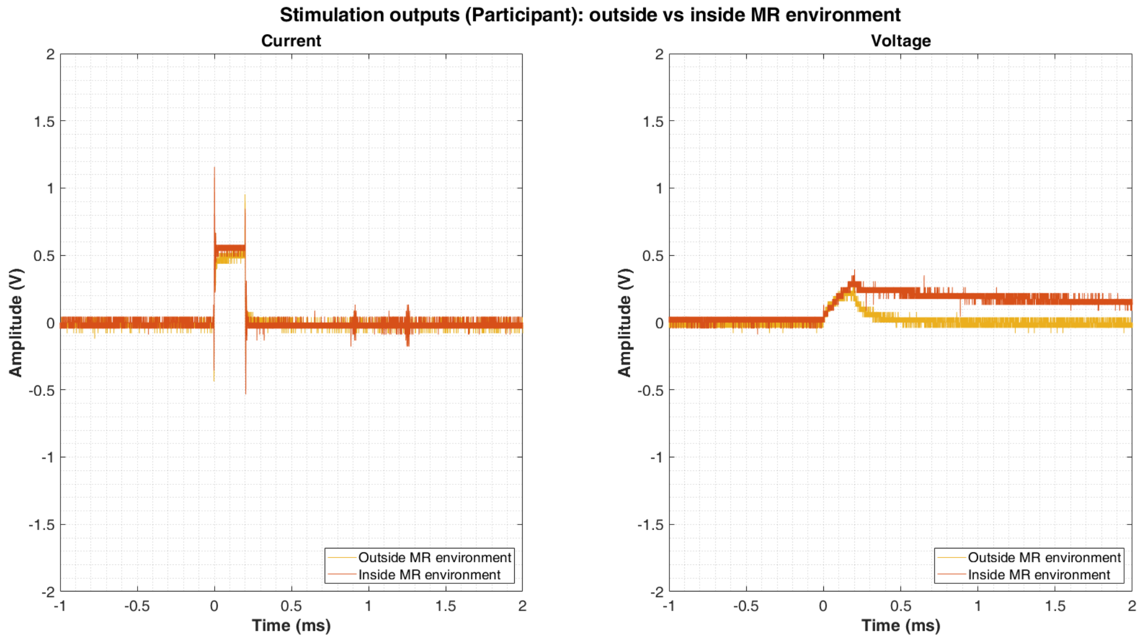


Figure S5.2 - Comparison of the current (left) and voltage (right) outputs measured inside (orange) and outside (yellow) the magnetic resonance (MR) environment for a single stimulus during participant acquisition. The overlapping waveforms highlight their similarity, indicating that the MR environment did not influence the performance of the stimulation device. Stimulation consists of positive rectangular pulses of 0.2 ms at a frequency of 100 Hz. Please note a slight disparity in stimulus intensity: the current intensity outside the MR environment was 1 mA, while inside the MR environment it was 1.10 mA. Differences in voltage intensity can be attributed to current intensities and factors associated with skin resistance (e.g. hydration, exfoliation, etc).

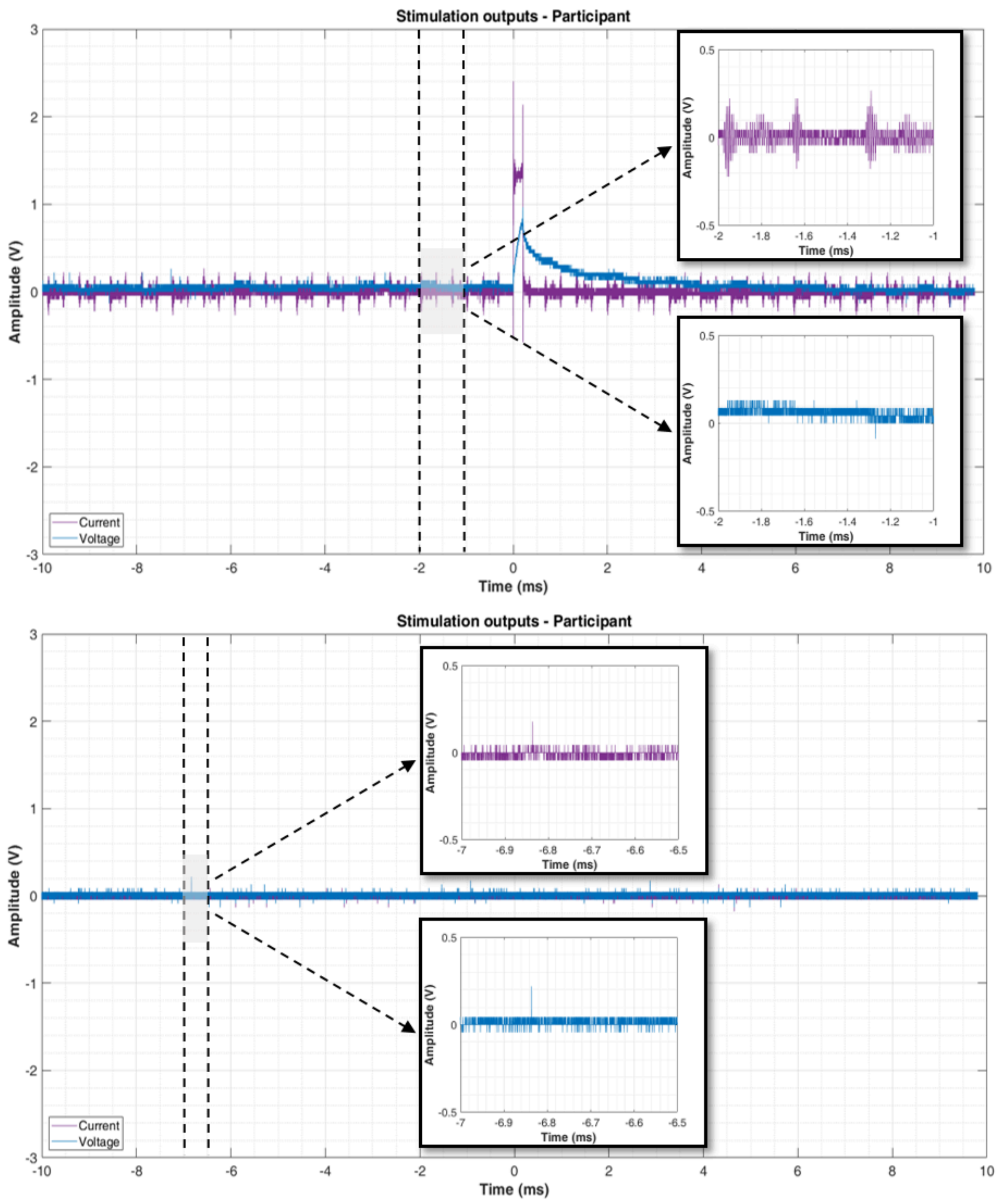


Figure S5.3 - Comparison of the noise levels at the current (purple) and voltage (blue) outputs measured inside the magnetic resonance (MR) environment during a functional acquisition with a participant. In the top image is represented one of the worst-case scenarios in terms of noise for a single stimulation pulse (Device operating); in the bottom image is represented the scenario where device is ON and the fMRI sequence is running (Device not operating). The maximum noise values for current and voltage in the Device operating condition are 0.27 V and 0.26 V, which corresponds to 0.53 mA and 10.54 V, respectively. In the Device not operating condition, the maximum noise values for current and voltage are 0.18 V and 0.22 V, corresponding to 0.36 mA and 8.78 V, respectively. The maximum noise values for the Device operating condition, considering a worst-case scenario, are approximately 1.5 times higher for current and 1.2 times higher for voltage compared to the Device not operating condition. Notably, owing to the design of our device, the relatively insignificant noise in the voltage waveform profile does not exert any discernible impact on the stimulus (output signal) experienced by the participant.
